# Tandem mobilization of anti-phage defenses alongside SCC*mec* cassettes

**DOI:** 10.1101/2023.03.17.533233

**Authors:** Motaher Hossain, Barbaros Aslan, Asma Hatoum-Aslan

## Abstract

Bacterial viruses (phages) and the immune systems targeted against them significantly impact bacterial survival, evolution, and the emergence of pathogenic strains. While recent research has made spectacular strides towards discovering and validating new defenses in a few model organisms^1-3^, the inventory of immune systems in clinically-relevant bacteria remains under-explored, and little is known about the mechanisms by which these systems horizontally spread. Such pathways not only impact the evolutionary trajectory of bacterial pathogens, but also threaten to undermine the effectiveness of phage-based therapeutics. Here, we investigate the battery of defenses in staphylococci, opportunistic pathogens that constitute leading causes of antibiotic-resistant infections. We show that these organisms harbor a variety of anti-phage defenses encoded within/near the infamous SCC (staphylococcal cassette chromosome) *mec* cassettes, mobile genomic islands that confer methicillin resistance. Importantly, we demonstrate that SCC*mec*-encoded recombinases mobilize not only SCC*mec*, but also tandem cassettes enriched with diverse defenses. Further, we show that phage infection potentiates cassette mobilization. Taken together, our findings reveal that beyond spreading antibiotic resistance, SCC*mec* cassettes play a central role in disseminating anti-phage defenses. This work underscores the urgent need for developing adjunctive treatments that target this pathway to save the burgeoning phage therapeutics from suffering the same fate as conventional antibiotics.

---

Bacterial viruses (phages) and the immune systems targeted against them significantly impact bacterial survival, evolution, and the emergence of pathogenic strains. While recent research has made spectacular strides towards discovering and validating new defenses in a few model organisms^1–3^, the inventory of immune systems in clinically-relevant bacteria remains underexplored, and little is known about the mechanisms by which these systems horizontally spread. Such pathways not only impact the evolutionary trajectory of bacterial pathogens, but also threaten to undermine the effectiveness of phage-based therapeutics. Here, we investigate the battery of defenses in staphylococci, opportunistic pathogens that constitute leading causes of antibiotic-resistant infections. We show that these organisms harbor a variety of anti-phage defenses encoded within/near the infamous SCC (staphylococcal cassette chromosome) *mec* cassettes, mobile genomic islands that confer methicillin resistance. Importantly, we demonstrate that SCC*mec*-encoded recombinases mobilize not only SCC*mec*, but also tandem cassettes enriched with diverse defenses. Further, we show that phage infection potentiates cassette mobilization. Taken together, our findings reveal that beyond spreading antibiotic resistance, SCC*mec* cassettes play a central role in disseminating anti-phage defenses. This work underscores the urgent need for developing adjunctive treatments that target this pathway to save the burgeoning phage therapeutics from suffering the same fate as conventional antibiotics.

Staphylococci are ubiquitous skin-dwelling bacteria that play critical roles in health and disease. Over 40 different human-associated *Staphylococcus* species have been identified^4, 5^, and the majority are considered skin commensals with neutral or even positive impacts^6^. However, two species in particular, *S. aureus* and *S. epidermidis*, have significant pathogenic potential—*S. aureus* causes a wide array of hospital-and community-acquired infections, including bacteremia, osteomyelitis, and skin and soft tissue infections^7^; and *S. epidermidis* is the most common cause of infections associated with implanted medical devices^8^.

Compounding the problem, *S. epidermidis* harbors a reservoir of fitness/virulence factors which can be horizontally transferred to *S. aureus*^5, 9^. These include genes responsible for methicillin resistance (*mecA/C*) encoded on SCC (staphylococcal cassette chromosome) *mec* cassettes^10, 11^. Methicillin-resistant *S. aureus* (MRSA) poses a serious threat to global public health^12, 13^ and the disease burden has only worsened following the COVID-19 pandemic^14^. Further, multi-drug resistant *S. epidermidis* strains constitute an emerging global threat^15^. Thus, in order to develop effective therapeutic alternatives, it is critical to understand the major pathways that control the horizontal transfer of fitness/virulence factors between these species.

Bacterial viruses (phages) and the immune systems targeted against them have profound and opposing impacts on bacterial survival, evolution, and horizontal gene exchange^16^. For instance, strictly lytic phages can kill their host within minutes of infection, and accordingly are being harnessed for therapeutic applications to eradicate staphylococcal infections^17–19^. In stark contrast, the lysogenic/temperate phages may integrate into the host chromosome and are known to boost pathogenic potential by transferring virulence factors or pathogenicity islands to the host^16^. In response to the constant pressure of phage predation, bacteria have evolved a variety of immune systems that counter these diverse effects. At least four such systems have been identified and functionally validated in *S. aureus* and *S. epidermidis* strains—restriction-modification (RM)^20^, CRISPR-Cas^21^, a mechanism of abortive infection (i.e. programmed cell death) facilitated by the serine-threonine kinase Stk2^22^, and a unique innate immune system mediated by the nuclease-helicase Nhi^23^. These systems antagonize lytic and lysogenic phages alike, and therefore have the capacity to not only curb pathogenic potential, but also compromise the effectiveness of phage-based therapeutics. Although significant headway has been made in recent years towards identifying and functionally validating the diverse immune repertoire in a few model organisms^1–3, 24, 25^, the full battery of anti-phage defenses in staphylococci has not been systematically explored. Additionally, little is known about the predominant pathways by which these systems horizontally spread.

To shed light on these issues, we first examined the distribution and localization of homologs for all known anti-phage defenses in RefSeq collections of *S. epidermidis* (*n*=89) and *S. aureus* strains (*n*=982). This was accomplished by programmatically invoking MacSyFinder^26^ for each genome using the Hidden Markov Models and system definitions library of DefenseFinder^27^. The results revealed that staphylococci possess a diverse battery of defenses comprising at least forty distinct immune system types (Extended data Figure 1 and Extended data Table 1). Further, we noted that about half of the organisms in the dataset harbor 50% or more of their defenses within 300 genes downstream of *rlmH* (Fig. 1A). *rlmH*, also called *orfX*, is a core housekeeping gene downstream of which SCC*mec* cassettes are known to reside^28, 29^. These observations prompted us to hypothesize that SCC*mec* cassettes constitute a major vehicle through which staphylococci disseminate anti-phage defenses.

**Figure 1.**
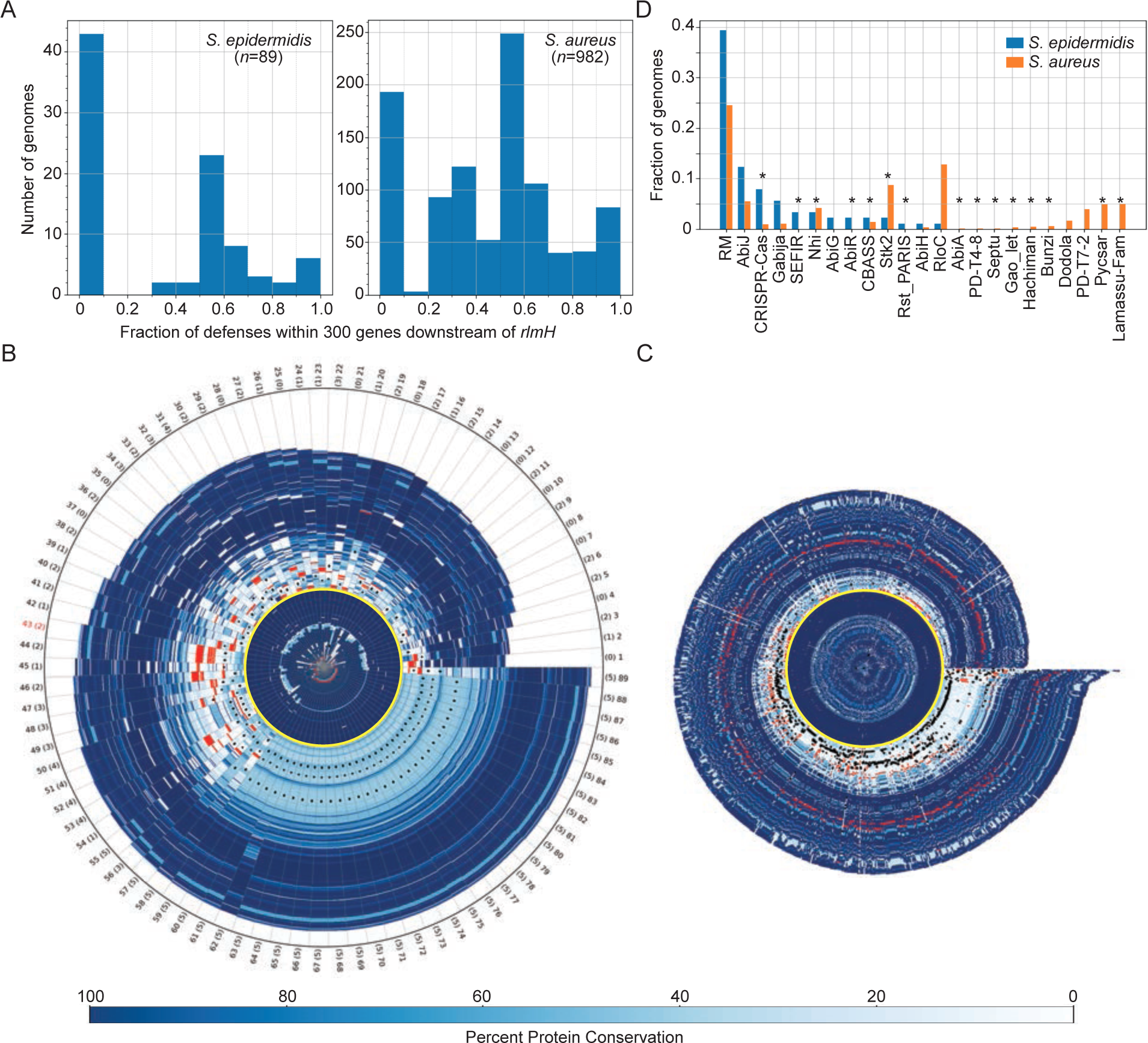
**SCC*mec* and adjacent accessory regions are rich with diverse defenses** (A) Histograms showing fractions of defenses encoded within 300 genes downstream of *rlmH* in sequenced *S. epidermidis* and *S. aureus* genomes. (B, C) Genome segments of *S. epidermidis* (B) and *S. aureus* (C) strains showing (from origin to tip) 70 proteins upstream of RlmH, variable numbers of accessory proteins downstream of RlmH, and an additional 70 proteins beyond the accessory/cassette region. RlmH (yellow ring), proteins with predicted defense functions (red bars), and positions of Ccr homologs (black dots) are highlighted. All other proteins are indicated in shades of blue that correspond to level of protein conservation across all genomes within each set (scale on bottom). The rightward boundary of the accessory region is defined as the occurrence of three consecutive proteins with over 95% conservation across all genomes within each set. Tip labels in (B) show genome number (1-89) and numbers of Ccr homologs detected (0-5, in parentheses). The tip label for *S. epidermidis* RP62a is shown in red. (D) A plot showing the types and distributions of anti-phage defenses encoded in the accessory regions of *S. epidermidis* and *S. aureus* genomes.

To test this idea, we performed a more detailed analysis of the proteins proximal to RlmH—all proteins encoded 200 genes upstream and 500 genes downstream of *rlmH* were analyzed for their identities, levels of conservation, and predicted defense functions. The results are depicted in a polar graph showing the protein content for the collections of *S. epidermidis* and *S. aureus* genome segments as spokes on a wheel (Fig. 1B and C, respectively and Extended data tables 2 and 3). The plots clearly show that while RlmH (yellow ring around the origin) is preceded by highly-conserved (*i.e.* core) proteins upstream (dark blue bars in center), it is followed by a sharp transition into a region of poorly-conserved (*i.e.* accessory) proteins downstream (light blue bars). These accessory regions are flanked by another stretch of highly-conserved proteins further downstream (dark blue periphery). If we define the upstream boundary of the ‘accessory region’ as RlmH, and the downstream boundary as the occurrence of three consecutive proteins that exceed 95% conservation across all genomes, then the accessory region lengths exhibit a range (5^th^ to 95^th^ percentile) spanning 54 to 136 proteins for *S. epidermidis* and 16 to 83 proteins for *S. aureus*. Strikingly, genes that encode known defense systems (red bars) are almost exclusively concentrated in the accessory regions. Also, the presence of between one and five putative Ccr recombinases (black dots)—the enzymes that mobilize SCC*mec* cassettes^30, 31^—suggest that the majority of genomes likely harbor at least one SCC*mec* cassette. There are at least 23 different defense types within the accessory regions, and interestingly, the majority of these (*n*=15) are located exclusively in this SCC*mec* region, including CRISPR-Cas, Stk2, and Nhi (Fig. 1D and Extended data Table 1). Taken together, these observations suggest that beyond carrying antibiotic resistance, SCC*mec* cassettes may host a variety of anti-phage defenses. However, SCC*mec* cassettes are typically 24-68 kilobases (kb) in length and contain ∼20-100 genes, respectively^28, 29^, and many of the accessory regions downstream of *rlmH* extend well beyond these lengths (Fig. 1C and D). Therefore, it is unlikely that all defenses are encoded within the bounds of the SCC*mec* cassettes.

To investigate further, we examined more closely *S. epidermidis* RP62a as a representative of the sequenced set (number 43 in Fig. 1B). This organism harbors a Type II SCC*mec* cassette (∼48 kb in length) that encodes ∼50 proteins, including the CcrA and CcrB recombinases^11^. These enzymes bind 18-nucleotide attachment (*att*) sites flanking SCC*mec*^32^ and catalyze cassette excision and circularization as prerequisite steps for the inter-*genus* horizontal transfer of the entire cassette^30, 31^. Our manual inspection of the RP62a SCC*mec* region revealed that only one known defense (*stk2*) lies within the bounds of the SCC*mec* cassette (Fig. 2A). However there appear to be at least three additional CcrAB *att* consensus sequences between SCC*mec* and the remaining defenses (Fig. 2A and B). These observations hinted at the compelling possibility that defenses encoded proximal to SCC*mec* may also be mobilized by CcrAB as separate/independent cassettes. To test this, we introduced the *ccrAB* operon from *S. epidermidis* RP62a under its native promoter into a multi-copy plasmid and transferred the plasmid (called pSepiCcrAB) into *S. epidermidis* RP62a. Cells were then grown to mid-log, and their DNA was extracted and subjected to PCR using a set of primer pairs specifically designed to detect excision and circularization of SCC*mec* and the putative defense-containing cassettes (Fig. 2A and Supplementary Table 1). The results revealed that overexpression of *ccrAB* indeed stimulates excision and circularization of not only SCC*mec*, but also two adjacent independent cassettes containing the Nhi, RM, and CRISPR-Cas systems (Fig. 2 C-E). The Nhi-RM and CRISPR-Cas cassettes are ∼17 kb and ∼26 kb in length, respectively. The cassettes can be excised independently, in pairs, or all three can be found linked together (Extended data Figure 2 A and B). Strikingly, the excision of all three cassettes could be readily detected by conventional PCR even without CcrAB overexpression (Fig. 2 F and G).

**Figure 2.**
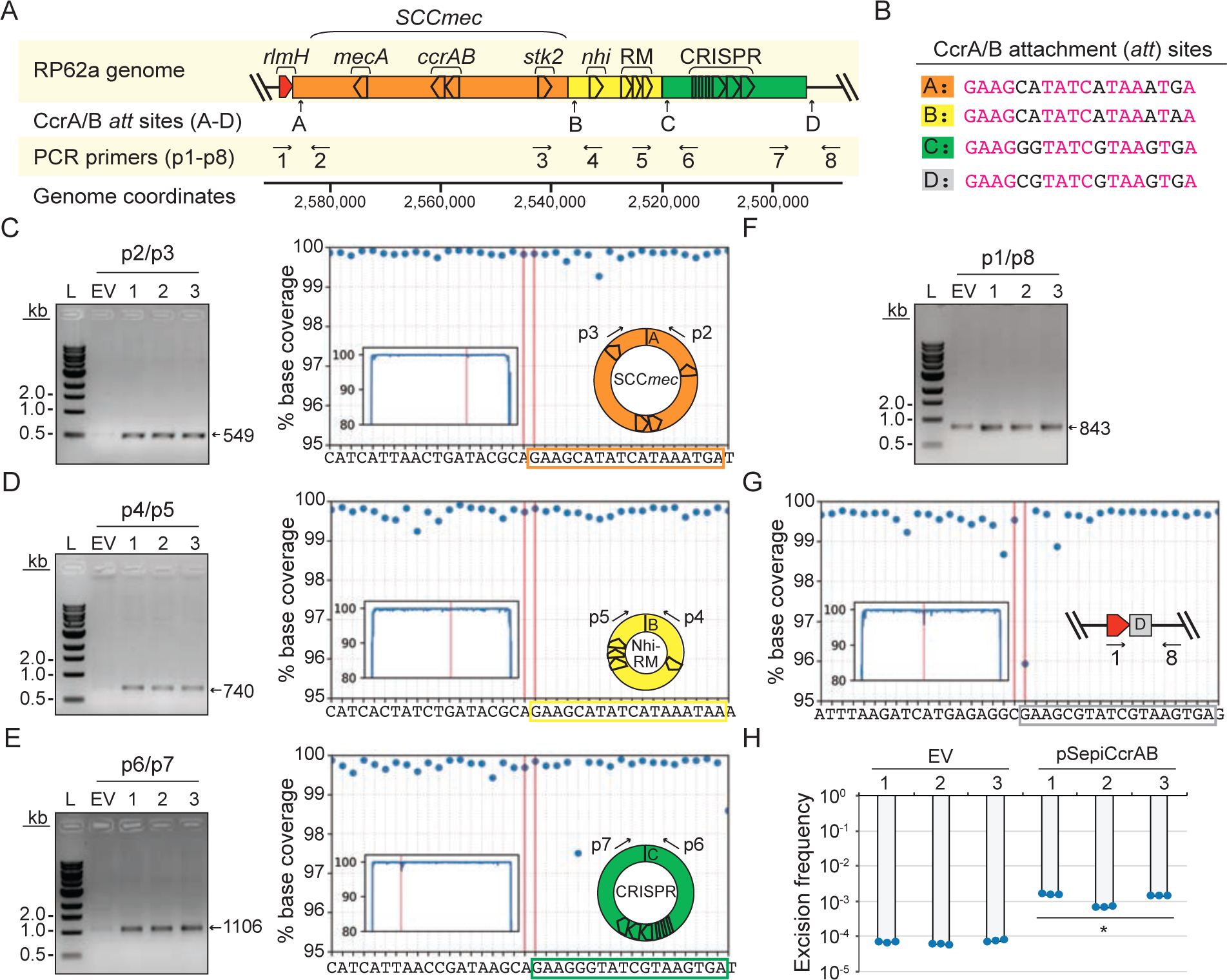
**Overexpression of *ccrAB* promotes mobilization of defense-enriched cassettes** (A) Illustration of *S. epidermidis* RP62a genomic region encoding SCC*mec* and known defenses (*stk2*, *nhi*, RM, and CRISPR-Cas). Positions of putative CcrAB attachment (*att*) sites (A-D) and PCR primers (p1-p8) are shown. (B) CcrAB *att* sites A-D are shown with identical nucleotides in magenta. (C-E) PCR products amplified from circularized individual cassettes (SCC*mec*, Nhi-RM, and CRISPR, respectively) resolved on agarose gels (left) and confirmed via Illumina sequencing (right). (F, G) PCR products amplified from the new junction created by excision of all three cassettes resolved on an agarose gel (F) and confirmed via Illumina sequencing (G) For C-G, DNA was extracted from three independent transformants of *S.epidermidis* RP62a-pSepiCcrAB (1-3) or cells harboring the empty vector (EV) and used as templates for PCR reactions. Indicated PCR primers were used to amplify new junctions resulting from circularization/excision of cassettes. Illumina sequencing reads (from one representative PCR product) were mapped back to the expected product sequence and the fraction of reads covering each position is shown. Insets show reads mapped to the full product length and main plots zoom into the regions flanking circle/excision junctions. The two vertical lines mark the precise boundaries of the new junctions generated from cassette circularization/excision. (H) Excision frequencies of all cassettes in *S. epidermidis* RP62a cells harboring pSepiCcrAB or the empty vector in three independently-generated transformants (1-3) as measured by qPCR. Data shown represents an average of triplicate measurements (±S.D.). A two-tailed t-test was performed to determine significance and * indicates p < 0.05.

We also examined the SCC*mec* regions of two additional clinical isolates which harbor Type III CRISPR-Cas systems—*S. aureus* MSHR1132, a community-associated MRSA strain recently re-classified as *S. argenteus*^33, 34^, and *S. aureus* ST398 08BA02176, a livestock-associated strain recovered from a human surgical site infection^35^. MSHR1132 possesses a Type IVa SCC*mec* cassette that encodes the CcrAB recombinases, and the Type III CRISPR-Cas system appears to be located downstream of the cassette flanked by additional *att* sites (Extended data Figure 3 A and B). To test if CcrAB overexpression promotes excision/circularization of these tandem cassettes, the *ccrAB* operon with its native promoter was inserted into a multicopy plasmid to create pSarCcrAB, the plasmid was introduced back into the host, and DNA extracts were assayed for evidence of cassette excision/circularization using conventional PCR. The results showed that tandem cassettes are indeed generated, but unlike RP62a, cassette circularization in MSHR1132 can be detected even in the absence of pSarCcrAB (Extended data Figure 3 C and D, EV lane). Accordingly, excision of both cassettes was also detected without *ccrAB* overexpression (Extended data Figure 3 E and F). In lieu of CcrAB, some SCC*mec* cassettes are mobilized by a single serine recombinase, CcrC, and we wondered whether its overexpression stimulates defense mobilization. To test this, we examined *S. aureus* ST398 08BA02176, which harbors a Type III CRISPR-Cas system within the bounds of a Type V SCC*mec* cassette (Extended data Figure 4 A and B)^35^. We created pSauCcrC (which bears *ccrC* under its native promoter), introduced the plasmid into ST398, and assayed DNA extracts for cassette mobilization via PCR amplification. We found that similarly to *ccrAB* in *S. epidermidis* RP62a, *ccrC* overexpression in *S. aureus* ST398 stimulates excision and circularization of the CRISPR-containing SCC*mec* cassette (Extended data Figure 4 C and D).

**Figure 3.**
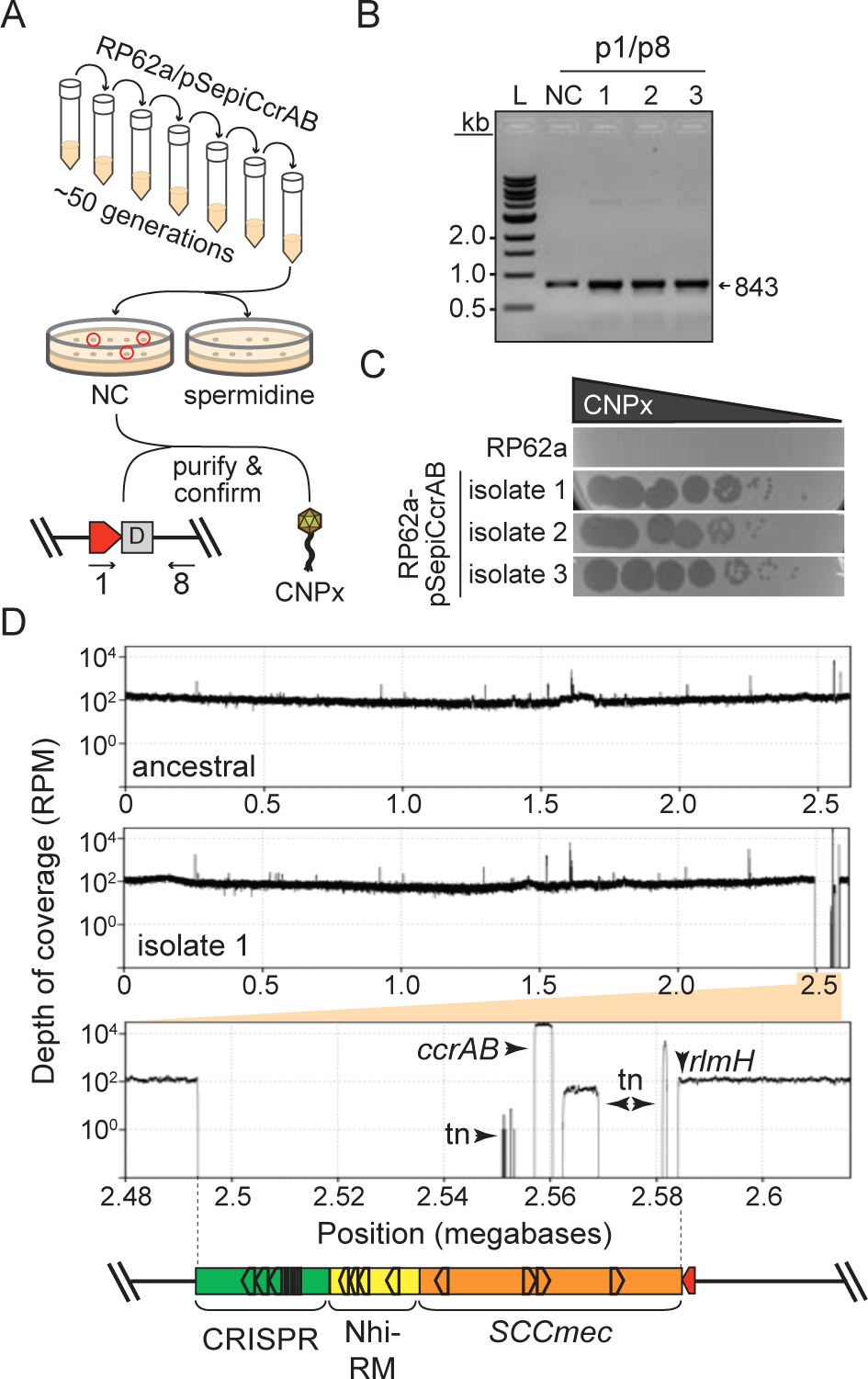
**CcrAB-mediated cassette excision is restricted to proximal genomic loci** (A) Illustration of the directed evolution approach used to screen for colonies that have lost all cassettes. (B) PCR amplicons derived from the new junction created by excision of all three cassettes in the *S. epidermidis* RP62a ancestral strain (NC) and three independently-generated evolved 1′cassette isolates. PCR products were resolved on an agarose gel. (C) A phage challenge assay is shown in which 10-fold dilutions of phage CNPx were spotted atop lawns of the *S. epidermidis* RP62a ancestral cells, or three independently-generated evolved 1′cassette isolates. (D) *S. epidermidis* RP62a ancestral and evolved 1′cassette strains were sequenced via Illumina, the reads for the ancestral strain were assembled, and the reads from the ancestral (top) and a representative evolved 1′cassette isolate (middle) were mapped back onto the wild-type/ancestral assembly. The plots show depth of coverage in reads per million (RPM) across the genomes. The bottom plot shows a close-up of the sole deleted genomic segment in the evolved 1′cassette isolate. Arrows indicate positions where read coverage originates from the plasmid-encoded *ccrAB* operon and transposable elements (tn) represented in other regions of the genome.

**Figure 4.**
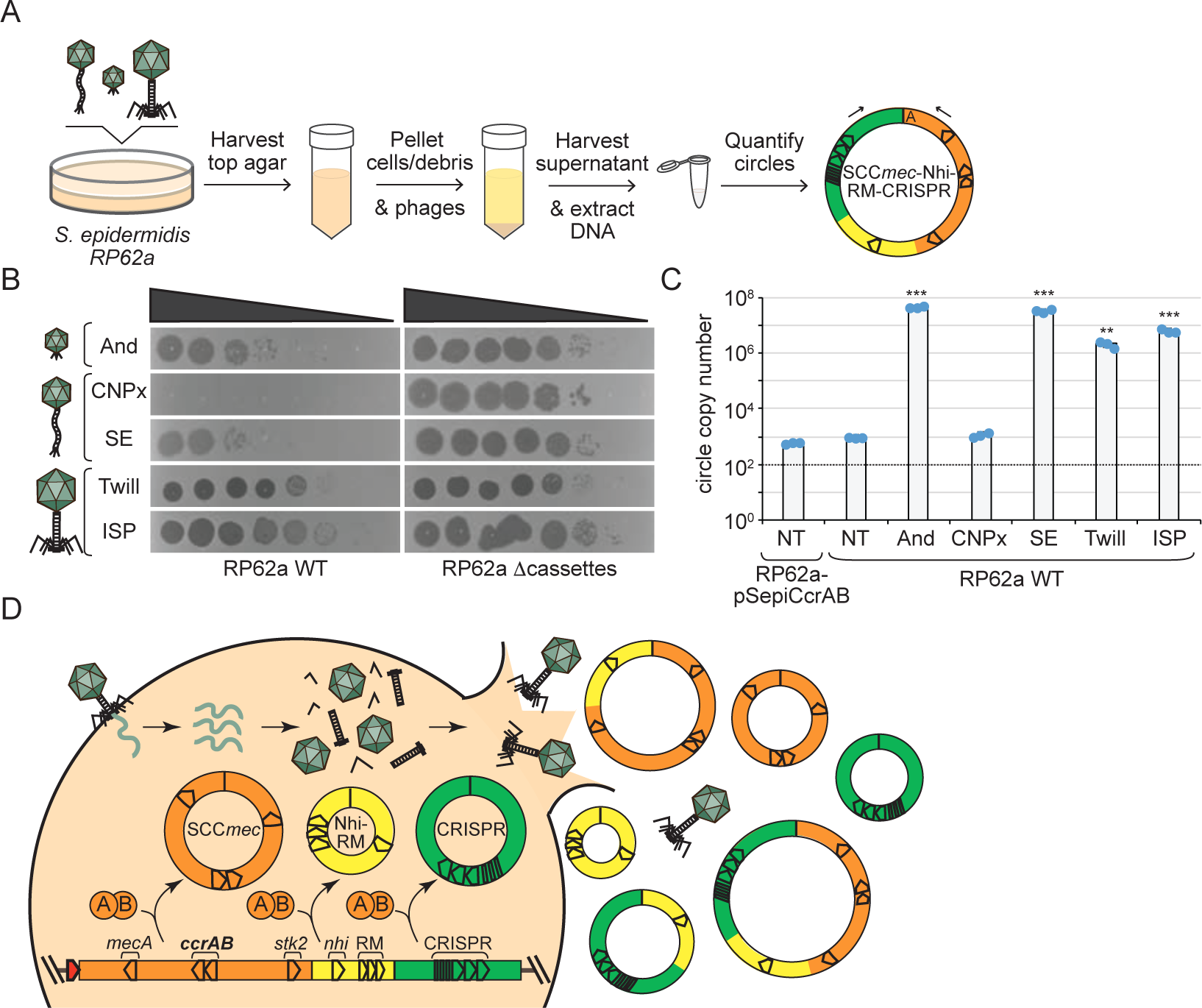
**Phage infection potentiates cassette release.** (A) Illustration of the assay used to quantify cassette release following challenge with phage. (B) Ten-fold dilutions of diverse phages spotted atop lawns of *S. epidermidis* RP62a wild-type (WT) and an evolved isolate that has lost SCC*mec* and tandem cassettes (1′cassettes). Abbreviated phage names are as follows--And, Andhra; SE, Southeast; Twill, Twillingate. (C) Numbers of circularized cassettes released from indicated *S. epidermidis* strains following phage challenge as measured by qPCR. Shown is an average of triplicate measurements (±S.D.) as a representative of three independent trials. NT, no treatment. A two-tailed t-test was performed to determine significance, ** indicates p < 0.005 and *** indicates p < 0.005. (D) Proposed mechanism for cassette release following phage infection.

Given that cassette excision occurs in the absence of Ccr overexpression, we sought to quantify the baseline excision frequencies in representative *S. epidermidis* and *S. aureus* strains. To do so, we used quantitative PCR (qPCR) to determine, in a given genomic DNA sample, the fraction of genome copies that have lost all cassettes. The results showed that in *S. epidermidis* RP62a, between one and six genomes per 100,000 copies exhibit spontaneous loss of all cassettes, and *ccrAB* overexpression causes over 10-fold increase in this excision frequency (Fig. 2H). A similar assay conducted for *S. aureus* ST398 revealed a baseline excision frequency of 10-fold lower (∼1 x 10^-6^), and this value similarly increased by an order of magnitude in the presence of *ccrC* overexpression (Extended data Figure 4E). Drawing on these collective observations—particularly the facts that (1) cassettes are found to be excised in all possible combinations, and (2) overexpression of Ccr recombinases that reside within SCCmec increases excision frequencies by more than 10-fold—we arrived at one of our pivotal conclusions: Ccr recombinases drive the mobilization of diverse defenses encoded both within the SCC*mec* cassette and just outside (i.e. proximal) of its boundaries in separate tandem cassettes.

In light of the above, we next wondered whether these enzymes can also cause excision and circularization of distal genomic loci. To investigate this possibility, we first took a bioinformatics approach to identify all *att* consensus sites in the sequenced *S. epidermidis* collection and assess their localization. This analysis identified 1,326 putative *att* sites across the 89 *S. epidermidis* genomes, and remarkably, a significant fraction of sites (47%) resides outside the accessory region downstream of *rlmH* (Extended data Figure 5A and Extended data Table 4). Moreover, the genomes harbor between one and six pairs of distal *att* sites within 150 kb (maximum) of each other on the same strand, thus demarcating putative distal cassettes (Extended data Figure 5A, red dots). Notably, sequence logos built from proximal and distal *att* motifs are absolutely conserved at positions 1, 2, 7, 8, and 13 (Extended data Figure 5B). These observations prompted us to investigate the extent to which CcrAB may mobilize additional genomic loci flanked by *att* consensus sites.

To test this, we examined more closely *S. epidermidis* RP62a, which harbors 14 putative *att* sites. We were surprised to find that eight of these sites exist within the accessory region, and a subset demarcate two additional putative cassettes that lie directly downstream of the CRISPR-containing cassette (Extended data Figure 5C). Further, six additional sites lie distal to SCC*mec*, and three of these demarcate two putative distal cassettes. To determine whether CcrAB can cause loss of these additional proximal/distal genomic loci, we used directed evolution to generate mutant strains that have lost all possible cassettes via long-term overexpression of *ccrAB*. Briefly, RP62a strains bearing pSepiCcrAB were passaged over ∼50 generations, and colonies were screened for the loss of resistance to spermine, which is conferred by *speG* within the Nhi-RM cassette (Fig. 3A). Colonies that exhibited sensitivity to spermine were purified and confirmed for loss of all cassettes downstream of *rlmH* by PCR amplification (Fig. 3B). In addition, to rule out the possibility that cassettes might still be present in the cell, perhaps in an alternative location or in a circularized episomal form, we challenged the spermine-sensitive isolates with phage CNPx, which is targeted independently by defenses encoded within each of the three cassettes: Stk2^22^, Nhi^23^, and CRISPR-Cas^36^. As expected, while CNPx cannot form plaques (i.e. zones of bacterial growth inhibition) on the ancestral RP62a strain, it forms millions of plaques on three independently-generated evolved isolates (Fig. 3C), thus confirming the complete loss of all three cassettes. To determine the extent to which these mutants have lost additional genomic loci, we purified genomic DNA from the isolates, subjected the DNA to Illumina sequencing, and mapped the sequencing reads back to an assembly of the RP62a ancestral genome. The results showed that contrary to our expectations, CcrAB-mediated genomic loss in RP62a is restricted to SCC*mec* and the two proximal defense-containing cassettes (Fig. 3D and Extended data Figure 6).

Finally, in light of our findings that tandem defense-enriched cassettes exist in an equilibrium of excised and integrated states in the absence of external stimulation (Fig. 2H and Extended data Figure 4E), we reasoned that phage infection likely potentiates cassette dissemination. To test this, we quantified the amounts of circularized cassettes released from cells following challenge with a panel of diverse phages (Fig. 4A). The phages exhibit a range of sensitivities to defenses encoded within the cassettes—from completely resistant (e.g. ISP) to partially or fully sensitive (CNPx) (Fig. 4B). The results showed that diverse phages indeed cause a striking stimulation of cassette release (between 10^3^ – 10^5^-fold) (Fig. 4C). Notably, neither CNPx infection nor CcrAB overexpression caused an increase in the number of extracellular cassettes. These observations support a model in which SCC*mec* and tandem cassettes are released from cells during phage-induced lysis (Fig. 4D).

While functional analyses of SCC*mec* cassettes have historically focused on the cargo of antibiotic resistance genes they carry, other accessory elements (encoded in so-called ‘junkyard’ regions) have remained largely uncharacterized^28, 29^. Here, we show that beyond spreading antibiotic resistance, SCC*mec* cassettes play a central role in disseminating diverse anti-phage defenses encoded both within and outside the bounds of the SCC*mec* cassette (Figs. 1 and 2). These observations support the notion that SCC*mec* and surrounding regions likely host a treasure trove of new defenses and other fitness factors yet to be identified. Moreover, the ability of SCC*mec*-encoded recombinases to mobilize additional genomic segments blurs the definition of what precisely constitutes an SCC*mec* cassette. The prevailing view is that SCC*mec* cassettes are discrete mobile elements with well-defined boundaries^28, 29^; however, our observations hint at a more expansive view of these elements as contiguous collections of accessory genes flanked by Ccr *att* sites. Akin to the engines of locomotives, Ccr recombinases need only be present in a single cassette to facilitate movement of the rest. In agreement with this notion, some *S. epidermidis* and *S. aureus* strains have been found to harbor ACME (arginine catabolic mobile element) and COMER (copper and mercury resistance) cassettes flanked by *att* sites downstream of SCC*mec*^32, 37–39^. Similar to the defense-enriched cassettes described in this study, ACME and COMER elements lack their own recombinases and are mobilized by those encoded within SCC*mec*.

The evolution of such modularity in cassette architecture may confer multiple advantages in facilitating cassette transmission, reception, and retention. For instance, collections of cassettes have the benefit of maintaining high carrying capacity while also keeping open the option of being mobilized as smaller nested segments which are more amenable to acquisition by the next host. Indeed, staphylococci are notoriously difficult to transform, and the predominant pathway by which SCC*mec* cassettes are acquired remains debated--Some studies showed that phages play a role in packaging and transfer of smaller SCC*mec* types^40^ or SCC*mec*-encoded elements^41, 42^ via generalized transduction, one study demonstrated SCC*mec* transfer can occur via conjugation^43^, and a recent report showed that whole cassettes can be acquired at low frequencies via natural transformation^44^. In light of the strict packaging constraints of staphylococcal transducing phages, combined with the longer and variable lengths of SCC*mec* and associated cassettes, it is not surprising that these elements have been found to rely upon multiple modes of transportation. Moreover, from a risk management perspective, genomic segments traveling independently as smaller cassettes maximizes their potential to survive in transit while being challenged with RM, CRISPR-Cas, or other nucleic acid degrading defenses. Finally, harboring multiple copies of *att* sites throughout nested cassettes ensures that at least a subset maintains mobility should random mutagenesis render one or more sites unrecognizable.

Our observation that phage lysis causes the release of defense-enriched cassettes alongside SCC*mec* (Fig. 4) allows for an unsettling prediction—the very same mechanisms that mediate the spread of methicillin resistance are likely to compromise the long-term effectiveness of phage-based therapeutics. These findings underscore the need for the preemptive development of adjunctive therapeutic strategies that intervene with the mobilization and transfer of these cassettes. It is also imperative to identify the other locations where anti-phage defenses reside. Indeed, not all of the analyzed staphylococcal genomes maintain the bulk of their defenses within/near SCC*mec* cassettes (Fig. 1A). This observation raises the question—where are the other defenses located? A recent study showed that *S. aureus* pathogenicity islands (SaPIs) constitute hotspots for anti-phage defenses^45^. SaPIs are a family of short (<20 kb) mobile elements that parasitize specific helper phages to facilitate their own packaging and spread. Indeed, there has been an increasing awareness that diverse bacteria harbor an assortment of defenses within mobile elements (MGEs) including similar parasitic phage-like elements (also known as satellites)^25, 45, 46^, plasmids^47^, transposons^48^, integrative conjugative elements^49^, and integrated temperate phages (prophages)^25, 50, 51^. Such defenses are thought to play major roles in inter-MGE conflicts. Our ongoing work continues to investigate the localization of defenses across staphylococci and seeks to identify new immune systems. These efforts will not only shed light on the predominant pathways that mediate phage-host interactions but are also likely to enable the development of more effective and robust alternative therapeutics.

## Methods

### Computational analyses

The RefSeq collections of *S. epidermidis* and *S. aureus* genomes were downloaded in GenBank format on Nov. 21, 2021 and July 5, 2022, respectively. RlmH was located in each genome, and all proteins were oriented in the same direction such that RlmH can be read from left to right. To generate the polar plots in Fig. 1B and C, the sequences of the 200 proteins upstream and 500 proteins downstream of RlmH were extracted and stored in a matrix where the rows correspond to protein positions (0 to 700), and the columns to the genomes in the dataset. To determine conservation levels, all protein sequences in the matrix were processed as a unified dataset using a hierarchical clustering algorithm we implemented. The algorithm dynamically aggregates protein sequences into clusters based on their pairwise similarity using the concept of Levenshtein edit ratio^52^ and a threshold value of 0.95. To determine the defense-relatedness flag (binary) for each protein sequence element of the matrix, we used MacSyFinder^26^ and Hidden Markov Model (HMM) system definition library of DefenseFinder^27^ to scan each genome for the presence of complete (known) defense systems. Only the component genes of a fully detected system were flagged as defense-related in the protein sequence matrix. In the end, the resulting matrix comprises protein sequences associated with two attributes: their conservation level, represented as a percentage value between 0 and 100%, and a binary indicator (0 or 1) that denotes whether a given protein sequence corresponds to a defense-related gene, along with the name of the corresponding HMM model. The other attributes such as whether the protein sequence is RlmH or identified as a Ccr recombinase are determined through the presence of associated protein family (pfam) domains using hmmer 3.3.2^53^. For the *att* consensus site analysis (Extended data Figure 5), an *att* consensus was derived from 16 verified CcrAB *att* sites compiled from the literature^32^ and our own observations (Fig. 2B), and a position-specific scoring matrix (PSSM) was calculated. Next, *rlmH* was located in each of the 89 *S. epidermidis* genomes, genome nucleotide sequences were oriented such that *rlmH* can be read from left to right, and the *rlmH* translational start site was designated as position 1. Using the PSSM, every preprocessed genome sequence was searched for matching motif hits (in both forward and reverse complement directions). Hits were subsequently clustered into two categories--those falling inside the ‘accessory region’ (*i.e.* light blue region downstream of *rlmH* in Fig. 1 B), and those falling outside. For hits outside the accessory region, a one-dimensional clustering algorithm was used to detect motif instance pairs on the same strand that were close enough to demarcate a putative cassette (i.e. <150 kb in length).

### Bacterial strains and growth conditions

*S. epidermidis* RP62a^54^ and *S. argenteus* MSHR1132^33^ were grown in Brain Heart Infusion (BHI, BD Diagnostics). *S. aureus* RN4220^55^ and ST398^35^ were grown in Tryptic Soy Broth (TSB, BD Diagnostics). Growth media was supplemented with 10 mg/mL chloramphenicol (for selection of pC194-based plasmids) and 15 mg/mL neomycin (for *S. epidermidis* strains). All bacterial strains were grown at 37°C. Liquid cultures were propagated with agitation in an orbital shaker set to 180 rpm. Strains were routinely authenticated via PCR amplification and sequencing genomic regions unique to each strain.

### Constructing ccr overexpression plasmids and strains

Plasmids pSepiCcrAB, pSarCcrAB, and pSauCcrC were created to overexpress Ccr recombinases. These plasmids were created using the multicopy plasmid pC194^56^ as backbone and genomic DNA from *S. epidermidis* RP62a, *S. argenteus* MSHR1132, and *S. aureus* ST398, respectively, to amplify *ccr* inserts. Plasmids were assembled via two-piece Gibson Assembly^57^ using the primers listed in Supplementary Table 1. Following Gibson assembly, constructs were introduced into *S. aureus* RN4220 by electroporation. Electrocompetent cells were prepared as described by Monk and colleagues^58^. For transformation, competent cells were thawed on ice for 5 min and left at room temperature for another 5 min. Cells were then pelleted via centrifugation at 5,000 *x g* for 1 min. The pellet was resuspended in 50 μL of sterilized 10% glycerol containing 500 mM sucrose, and the dialyzed Gibson assembly mix was added into the cell suspension. The mixture was then transferred into a 2 mm electroporation cuvette (VWR) and pulsed at 21 kV/cm, 100W, and 25 mF with a GenePulser Xcell instrument (Bio-Rad). Cells were then allowed to recover in 1 mL of sterile TSB containing 500 mM sucrose at 37°C with agitation for 2 hours. Recovered cells (200 μL) were plated on TSA or BHI agar supplemented with appropriate antibiotics and incubated at 37°C. Transformants were recovered the following day and confirmed for the presence of the intended plasmid by PCR amplification of the junctions of the assembled plasmids and Sanger sequencing using primers MH070 and F016 (Supplementary Table 1). At least three transformants were confirmed, confirmed plasmids were purified from *S. aureus* RN4220 using the E.Z.N.A.® Plasmid DNA Mini Kit I (Omega Bio-Tek, Inc, GA, USA), and purified plasmids pSepiCcrAB, pSauCcrC, and pSarCcrAB, were transferred into the appropriate strain (RP62a, ST398, and MSHR1132, respectively).

### Detecting cassette circle and excision junctions using conventional PCR

Amplification of circle and excision junctions was performed in 25 μL PCR reactions containing 25-100 ng of DNA template (plasmid miniprep), primers listed in Supplementary Table 1, and Phusion high fidelity DNA polymerase (NEB) according to the manufacturer’s instructions. PCR products were resolved on 1% agarose gels. For sequence confirmation, PCR products were purified using the E.Z.N.A.® Cycle Pure kit (Omega Bio-Tek, Inc, GA, USA), product concentrations were measured using the NanoDrop™ 2000 Spectrophotometer (Thermo Fisher Scientific), and products were submitted for Sanger sequencing (Eurofins Genomics, Louisville, KY) and/or Illumina sequencing (MiGS sequencing Center, Pittsburgh, PA).

### Preparing bacterial genomic DNA for Illumina sequencing and qPCR

Overnight cultures were diluted 1:100 in TSB/BHI and grown at 37°C until OD600 reached 1. Cultures (20 mL) were then transferred to 50 mL conical tubes and centrifuged at 5000 *x g* for 5 min at 4°C. Supernatants were discarded, washed once with 20 mL fresh TSB/BHI, and pellets were stored at −80°C. For DNA extraction, cell pellets were resuspended with 200 μL of sterile water and transferred into microtubes. Resuspended pellets were then incubated with lysostaphin (100 μg/mL) and MgCl_2_ (5 mM) at 37°C for 2 hours. The Wizard® Genomic DNA Purification Kit (Promega Corporation, WI, USA) was used to extract the genomic DNA according to the manufacturer’s instructions. Final DNA pellets were dissolved in 50-60 μL of prewarmed DNase-free water. DNA concentrations were measured using the NanoDrop™ 2000 Spectrophotometer (Thermo Fisher Scientific), and the samples were stored at 4°C for short-term use or at −20°C for long-term use.

### Constructing plasmids for qPCR standard curves

All PCR primers used for plasmid constructs are listed in Supplementary Table 1. Plasmids pSepiStdEx, pSauStdEx, and pSepiStdCirc were created to use as qPCR standards (Std) to measure numbers of cassette excision (Ex) and circle (Circ) junctions in *S. epidermidis* RP62a (pSepi) and *S. aureus* ST398 (pSau). All plasmids were constructed via two-piece Gibson assembly^57^ using PCR primers listed in Supplementary Table 1. Plasmid pC194^56^ was used as backbone, and genomic DNA for inserts as follows: The 894 bp insert for pSepiStdEx was amplified from the genomic DNA preparation of *S. epidermidis* RP62a Δ*cassette* strain (which contains the chromosomal junction formed upon the loss of SCCmec and tandem cassettes) using primers MH067 and MH068. The 960 bp insert for the pSepiStdCirc construct was amplified from a genomic DNA preparation of the *S. epidermidis* RP62a strain using primers MH109 and MH110. Similarly, the 858 bp insert of pSauStdEx was amplified from the genomic DNA of *S. aureus* ST398 using primers MH085 and MH086. All Gibson assembled constructs were transferred into *S. aureus* RN4220 and confirmed as outlined in the section above. Confirmed plasmids were purified, quantified, and used directly in qPCR assays to create standard curves.

### Measuring cassette excision frequencies and circle numbers using qPCR

All qPCR primers are listed in Supplementary Table 1. To determine cassette excision frequencies, primer pairs MH065/MH066 and MH089/MH090 were designed as controls to measure total genome copy numbers—these primers amplify the 5’-end of *rlmH* in *S. epidermidis* RP62a and *S. aureus* ST398, respectively. In addition, primer pairs MH081/MH084 and MH092/MH096 were designed to measure the numbers of genome copies that have lost all cassettes—these primers flank the cassette excision junctions in *S. epidermidis* RP62a and *S. aureus* ST398, respectively. To determine the number of released circularized cassettes, primers MH112/MH113 were designed to amplify the circle junction of all three cassettes in *S. epidermidis* RP62a. Each qPCR reaction (25 μL) consisted of 100 ng of genomic DNA as template, 0.4 nM control primers or 1 nM excision/circularization primers, and 1X PerfeCTa SYBR Green SuperMix (Quanta Biosciences). Separate sets of wells were prepared with appropriate standard reactions consisting of 10-fold dilutions of standard plasmids (pSepiStdEx, pSauStdEx, or pSepiStdCirc) containing 10^9^-10^2^ DNA molecules. The copy number of the standard plasmid molecules was calculated from the concentration and size of the standard plasmids using the URI Genomics & Sequencing Center online formula (https://cels.uri.edu/gsc/cndna.html). Each qPCR plate also included negative control wells containing nuclease-free H_2_O. The DNA templates were amplified using a CFX Connect Real-Time PCR Detection System (Bio-Rad) under the following conditions: one cycle of 95°C for 3 min; and 40 cycles of 95°C (10 sec) and 56.4°C (30 sec). At the end of the run, melt curves were generated to confirm homogenous products by exposing samples to a final temperature gradient of 65°C to 95°C. Following the reaction, standard curves were created using the CFX Maestro software (Bio-Rad), and a linear regression model was used to extrapolate product copy numbers. Excision frequencies represent the ratios of qPCR product copy numbers generated from the excision and control primer pairs.

### Constructing RP62a Δcassette strains via directed evolution

The RP62a/pSepiCcrAB strain was grown overnight and subcultured each day for seven consecutive nights (allowing ∼50 generations) in BHI (1:100) supplemented with neomycin and chloramphenicol at 37°C in a shaking incubator. The grown culture on the seventh day was serially diluted (10^0^-10^-7^) and 100 μL of the 10^-5^ dilution was spread onto BHI agar plates. The plates were incubated overnight at 37°C, and colonies were selected for further analysis. The colonies were resuspended into fresh BHI medium and then spotted onto both BHI agar and BHI agar supplemented with spermine (Acros Organics) at a final concentration of 0.9 mg/mL. Plates were incubated at 37°C overnight, and colonies that grew on BHI agar but not on BHI agar with spermine were selected. The selected colonies were confirmed for the loss of cassettes by amplifying the excision junction using PCR primers p1 and p8 and sequencing the products using Sanger sequencing. Colonies were also confirmed by plating with dilutions of phage CNPx according to the protocol described in the section on phage enumeration below.

### Mapping and assembling Illumina reads

For confirming cassette circle and excision junction PCR products via Illumina sequencing, Illumina paired-end reads were mapped onto the expected product sequence with Bowtie 2 v. 2.4.4. For analyzing genomic sequences of *S. epidermidis* RP62A-pSepiCcrAB ancestral and Δ*cassette* evolved isolates, paired-end reads for the ancestral genome were first assembled with SPAdes v. 3.15.3 using the reference sequence (NC_002976.3) as a template. Then, paired-end reads from three independently generated RP62A Δ*cassette* isolates were mapped back to the ancestral genome assembly using Bowtie 2 v. 2.4.4. Coverage was calculated from the resultant bam file (after sorting and indexing using samtools v. 1.13) using the "igvtools count" command, and normalized plots of coverage data were generated as a function of position, measured in reads per million.

### Phage propagation and enumeration

Phages CNPx^22^ (NC_031241), Southeast (OQ623150), Andhra^59^ (NC_047813), ISP^60^ (NC_047720), and Twillingate^61^ (MH321491) were propagated using *S. epidermidis* LM1680^62^ as host. To prepare the phage stocks, 1-5 purified phage plaques were combined into 500 μL TSB and vortexed for 30 sec. The suspension was then centrifuged at ∼15,000 *x g* for 2 min to pellet agar and cells, and the resulting phage lysate (i.e., supernatant) was passed through a 0.45 μm syringe filter. Next, the phage lysate was combined with overnight host culture (diluted 1:100) in 7 mL of Heart Infusion Agar (HIA) prepared at 0.3 x concentration and supplemented with 5 mM CaCl_2_. The phage-host mixture was poured onto a solid layer of Tryptic Soy Agar (TSA) supplemented with 5 mM CaCl_2_ and allowed to solidify for 10 minutes at room temperature. After overnight incubation at 37°C, the top agar layer was harvested and resuspended in 10 mL of fresh TSB. The suspension was vortexed for 5 min to release phages from the agar, followed by centrifugation at 10,000 *x g* for 10 min to remove agar and cell debris. The resulting concentrated phage lysates were passed through a 0.45 μm bottle filter to obtain a purified phage stock. Phage concentrations were determined by plating 10-fold dilutions of the phage suspension atop a lawn of cells using the double-agar overlay method as described by Cater and co-workers^59^. Phage stocks were stored at 4°C, and phages were routinely authenticated through PCR amplification and sequencing of genomic regions unique to each phage.

### Phage challenge in semi-solid agar and extraction of released DNA

The high titer phage lysates (≥10^10^ pfu/mL) propagated in *S. epidermidis* LM1680 were used to challenge *S. epidermidis* RP62a at a phage:bacteria ratio of 10:1 within soft agar overlays. Briefly, each phage lysate was combined with 300 μL of overnight host culture in 7 mL of Heart Infusion Agar (HIA) prepared at 0.3 x concentration and supplemented with 5 mM CaCl_2_. The phage-host mixture was poured onto a solid layer of Tryptic Soy Agar (TSA) supplemented with 5 mM CaCl_2_ and allowed to solidify for 10 min at room temperature. After 16-18 hours of incubation at 37°C, the top agar layer was harvested and resuspended into 10 mL of fresh TSB. The suspension was vortexed for 5 minutes, followed by centrifugation at 10,000 *x g* for 10 min to remove agar and cell debris. Finally, the resulting supernatant was passed through a 0.2 μm bottle filter to obtain a purified supernatant with no cellular debris. Lysates were further centrifuged at 22,000 *x g* and 4°C for 1 hour to pellet phages. The top portion of the supernatant from each tube was carefully collected in a 50 mL conical tube without disturbing the pellet. Next, 5 mL of the supernatant was mixed with an equal volume of Phenol:Chloroform:Isoamyl Alcohol (25:24:1) and vortexed for 1 minute. The mixture was subjected to centrifugation at 20,000 *x g* for 5 minutes at room temperature (RT), and the upper aqueous layer was transferred into a fresh tube. The aqueous phase was then mixed with sodium acetate (pH 5.2) to a final concentration of 200 mM, and two volumes of 100% ethanol was added. The tube was inverted 3-4 times and kept on ice for 10 minutes before centrifuging at 20,000 *x g* for 5 minutes at RT. The supernatant was gently decanted, and the pellet was washed with 3-5 mL of 75% ethanol, followed by centrifugation at 20,000 *x g* for 5 min at RT. After decanting the supernatant, the remaining liquid was carefully aspirated from the pellet, which was air-dried for 5-10 minutes. Finally, the pellet was resuspended in 100 μL of DNase-free dH_2_O, and its concentration was measured using NanoDrop™ 2000 Spectrophotometer (Thermo Fisher Scientific). DNA extracts were subjected to qPCR to quantify numbers of released circles as described in the section above.

### Statistical analyses

Graphed qPCR data represents the mean (±SD) of three replicates. Average values were analyzed in pairwise comparisons using two-tailed t-tests, and p-values < 0.05 were considered statistically significant. Sample sizes were empirically determined, and no outliers were observed or omitted.

## Data Availability

The RefSeq datasets for *S. epidermidis* and *S. aureus* genomes used in this study can be accessed with individual NCBI accession codes listed in Extended Data Tables 2 and 3, respectively. The raw Illumina sequencing reads generated in this study have been deposited in NCBI under Bioproject PRJNA945578.

## Code Availability

The custom code for analyzing RefSeq datasets can be made available upon request.

## Acknowledgments

This work was funded by NIH/NIAID [R01AI 173022-01]. A. H.-A. holds an Investigators in the Pathogenesis of Infectious Disease Award from the Burroughs Wellcome Fund. She is also supported by an NSF/MCB CAREER award [2054755], and funding from NIH/NIAID [R21AI156636-01].

## Author Contributions

M.H. designed and performed all of the wet-lab experiments, analyzed data, wrote methods, and reviewed and edited the manuscript; B.A. designed and performed all of the computational and bioinformatics analyses, analyzed data, wrote methods, and reviewed and edited the manuscript; and A.H-A. conceived of the study, acquired funding, supervised the work, analyzed data, and wrote the original draft of the manuscript. All authors approve of the authorship and content of the manuscript.

## Materials & Correspondence

Correspondence and requests for materials should be addressed to Asma Hatoum-Aslan at.

## Supplementary data

Supplementary Table 1. DNA oligonucleotides used for cloning and PCR in this study.

## Extended Data Tables

**Extended data Table 1.** Accompanies Extended data Fgure 1. Defenses in *Staphylococcus* genomes encoded within 300 genes downstream of *rlmH* and outside of that region

**Extended data Table 2.** Accompanies Figure 1B. Defenses in *S. epidermidis* genomes encoded within and outside of the accessory region downstream of *rlmH*

**Extended data Table 3.** Accompanies Figure 1C. Defenses in *S. aureus* genomes encoded within and outside the accessory region downstream of *rlmH*

**Extended data Table 4**. Accompanies Extended data Figure 5. Ccr *att* consensus sequences across *S. epidermidis* genomes within and outside of the accessory region downstream of *rlmH*.

## Extended Data Figures

**Extended data Figure 1.**
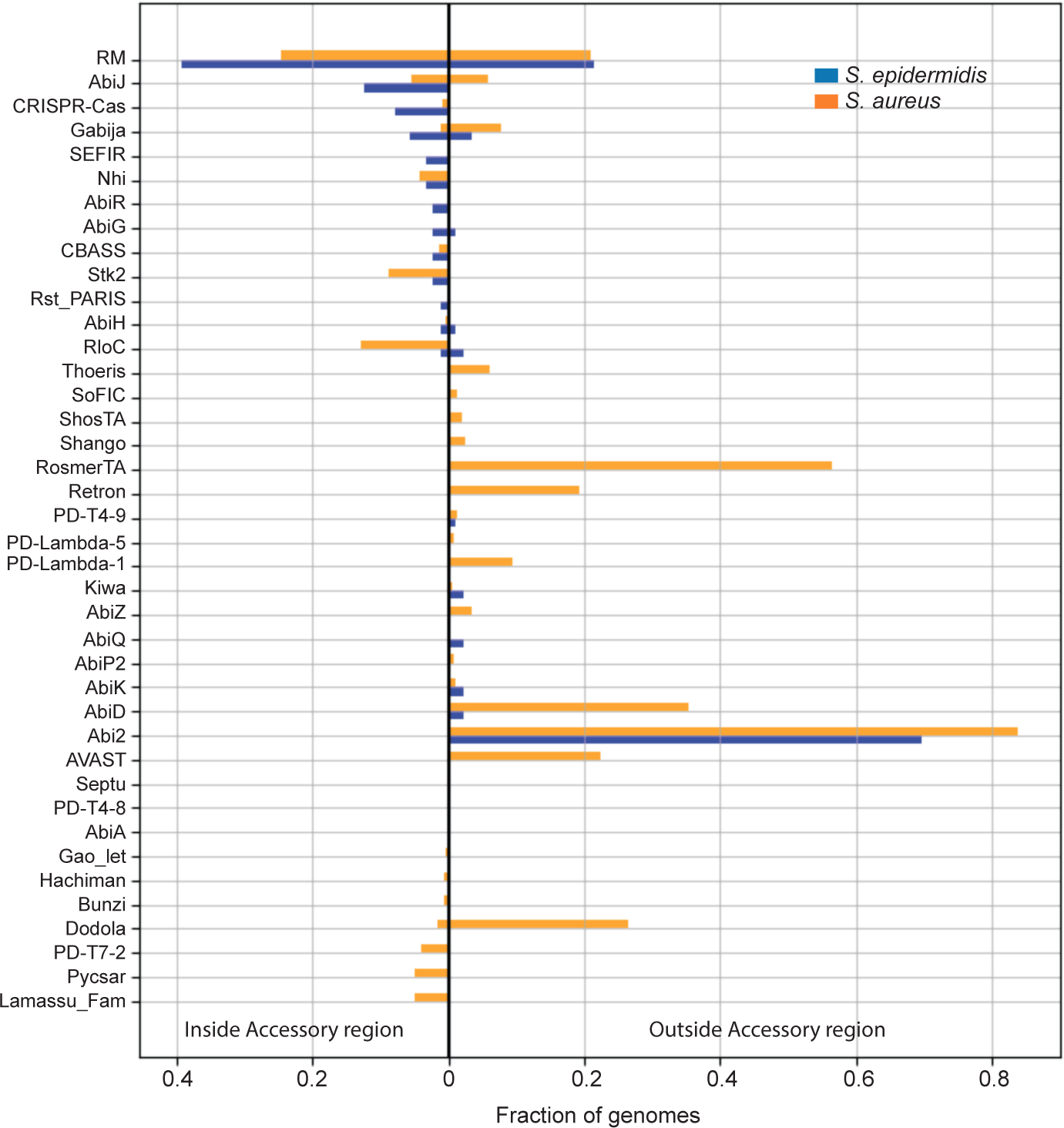
Accompanies Figure 1. Diversity and distribution of known defenses in *S. epidermidis* and *S. aureus* strains. The defenses are separated into two categories—those encoded within the accessory region downstream of *rlmH* (light blue regions in Fig. 1 B and C) and those encoded outside the accessory region.

**Extended data Figure 2.**
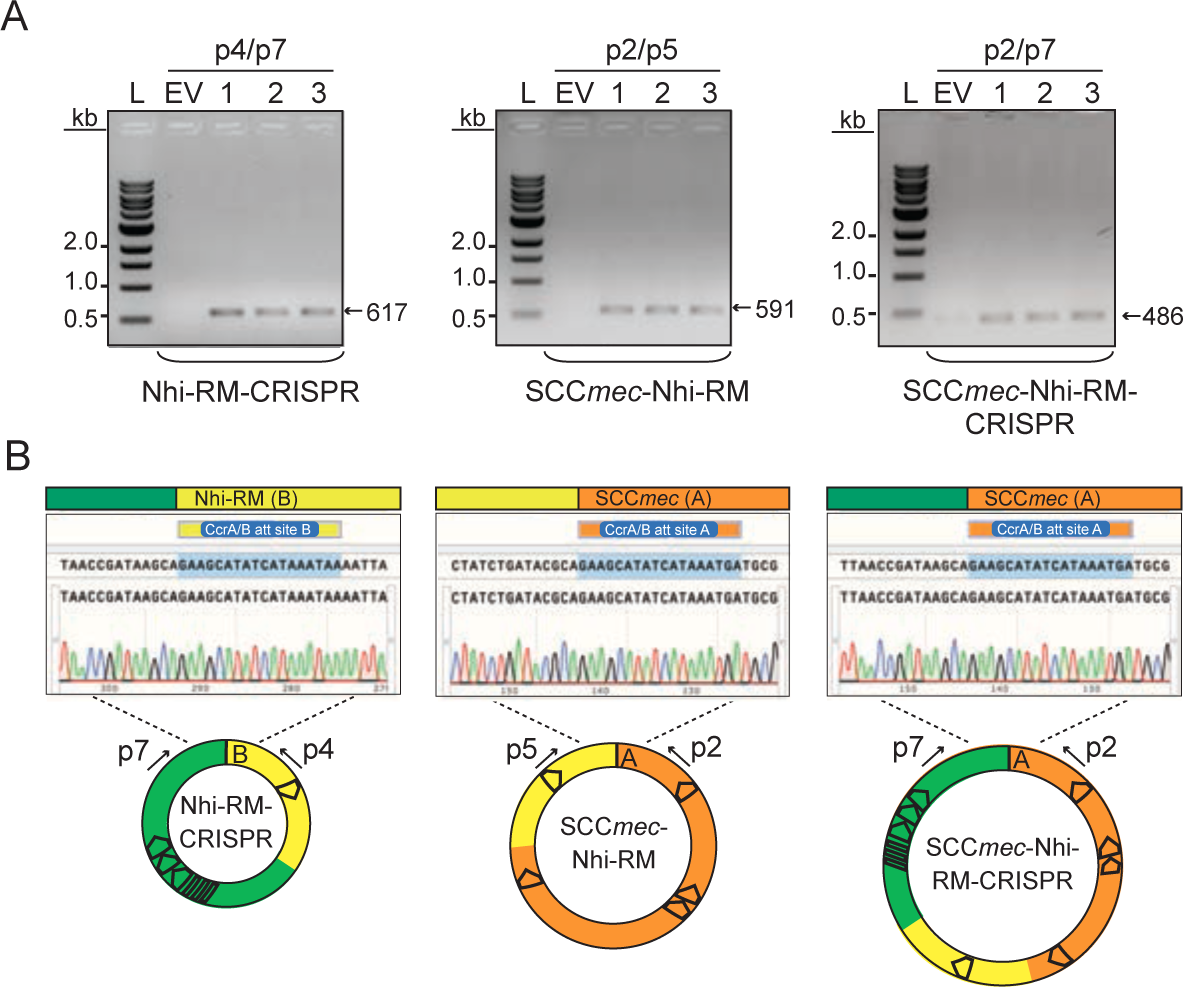
Accompanies Figure 2. Detection of composite cassettes in *S. epidermidis* RP62a. (A) PCR products amplified from composite circularized cassettes resolved on an agarose gel. DNA was extracted from three independent transformants of *S.epidermidis* RP62a-pSepiCcrAB (1-3) or cells harboring the empty vector (EV) and used as templates for PCR reactions. Indicated PCR primers were used to amplify new junctions resulting from circularization of cassettes. See Figure 2A for primer positions. (B) Sanger sequencing reads covering indicated circle junctions.

**Extended data Figure 3.**
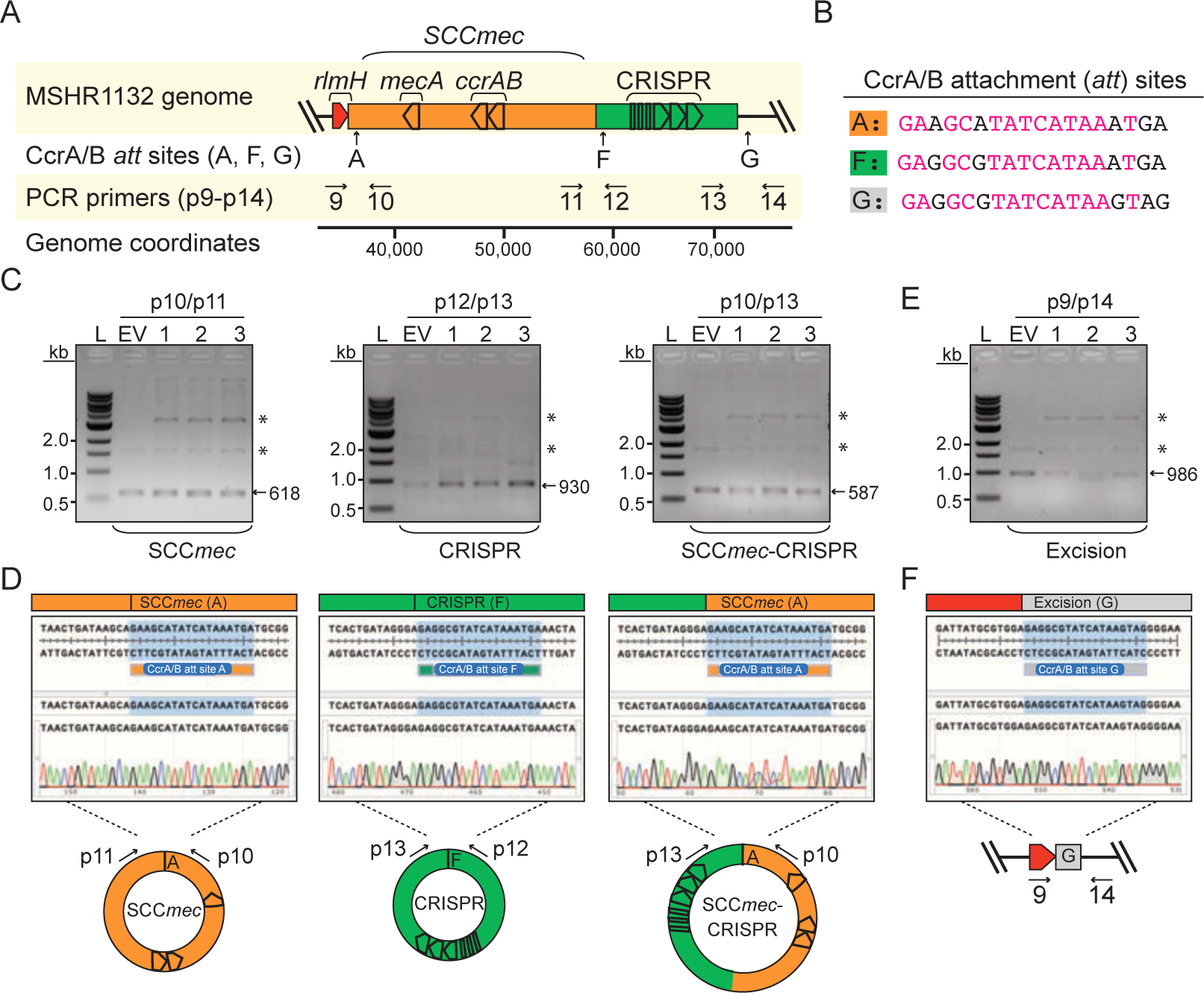
Accompanies Figure 2. Mobilization of an independent defense-containing cassette in *S. argenteus* MSHR1132. (A) Illustration of the genomic region encoding SCC*mec* and proximal CRISPR system. Positions of CcrAB attachment (*att*) sites (A, F and G) and PCR primers (p9-p14) are shown. (B) CcrAB *att* sites A, F and G are shown with identical nucleotides in magenta. (C, E) PCR products amplified from circularized cassettes (C) and the excision junction formed by loss of all cassettes (E) resolved on agarose gels. DNA was extracted from three independent transformants of *S. argenteus* MSHR1132-pSarCcrAB (1-3) or cells harboring the empty vector (EV) and used as templates for the PCR reactions. Indicated PCR primers were used to amplify new junctions resulting from circularization/excision of cassettes. Asterisks mark bands from the pSarCcrAB and EV plasmids in the DNA extract. (D, F) Sanger sequencing reads covering indicated circle/excision junctions.

**Extended data Figure 4.**
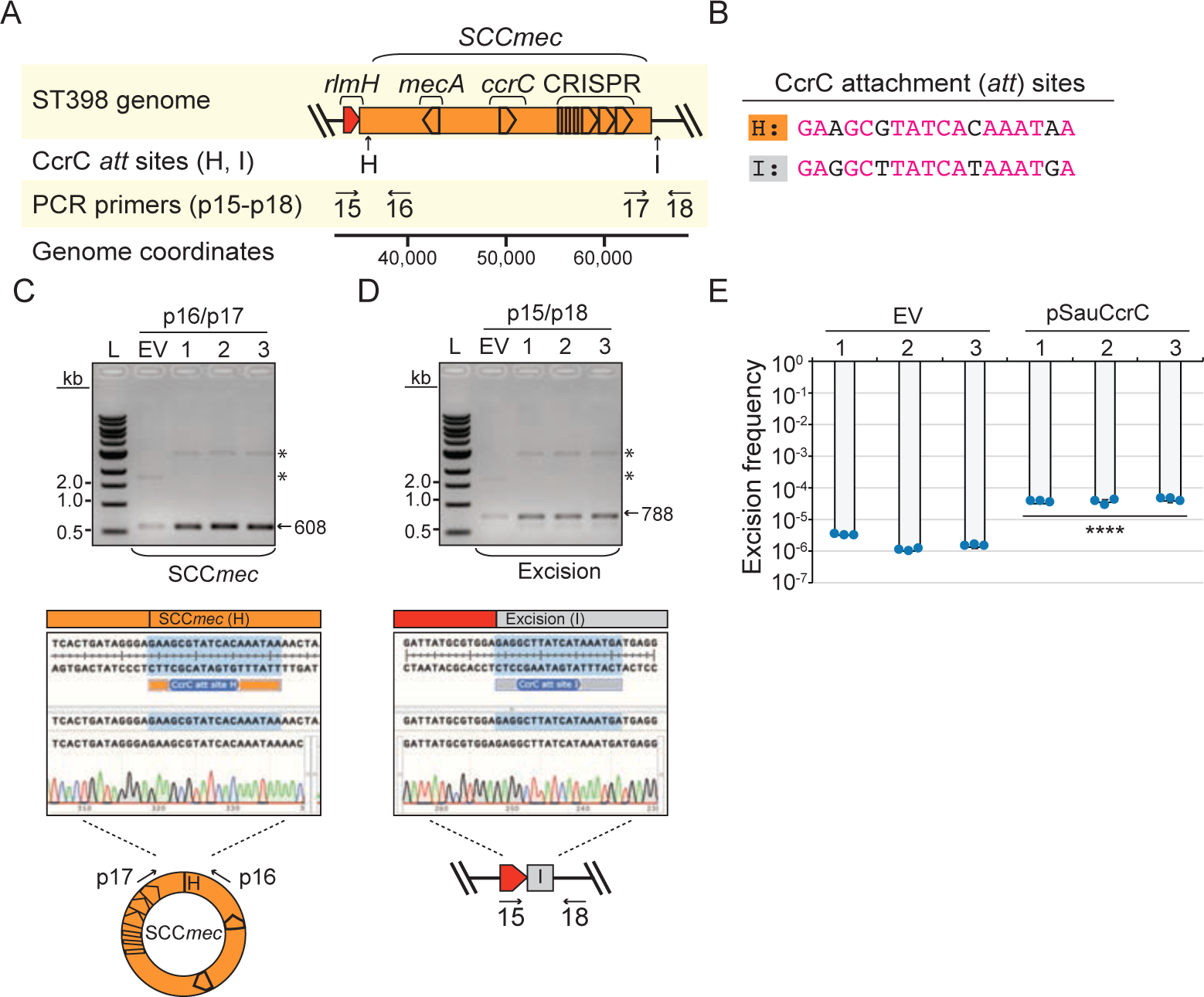
Accompanies Fig. 2. CcrC overexpression stimulates mobilization of an SCC*mec* cassette containing a CRISPR-Cas system in *S. aureus* ST398 08BA02176. (A) Illustration of the genomic region encoding SCC*mec* containing a CRISPR system. Positions of CcrC attachment (*att*) sites (H and I) and PCR primers (p15-p18) are shown. (B) CcrC *att* sites H and I are shown with identical nucleotides in magenta. (C, D) PCR products amplified from the circularized cassettes (C) and the excision junction formed by loss of the cassette (D) resolved on agarose gels (top). DNA was extracted from three independent transformants of *S. aureus* ST398-pSauCcrC (1-3) or cells harboring the empty vector (EV) and used as templates for the PCR reactions. Indicated PCR primers were used to amplify new junctions resulting from circularization/excision of cassettes. Asterisks mark bands from the pSauCcrC and EV plasmids in the DNA extract. Sanger sequencing reads covering indicated circle/excision junctions are also shown (bottom). (E) Excision frequencies of the cassette in *S. aureus* ST398 cells harboring pSauCcrC or the empty vector in three independently-generated transformants (1-3) as measured by qPCR. Data shown represent an average of triplicate measurements (±S.D.). A two-tailed t-test was performed to determine significance and **** indicates p < 0.00005.

**Extended Data Figure 5.**
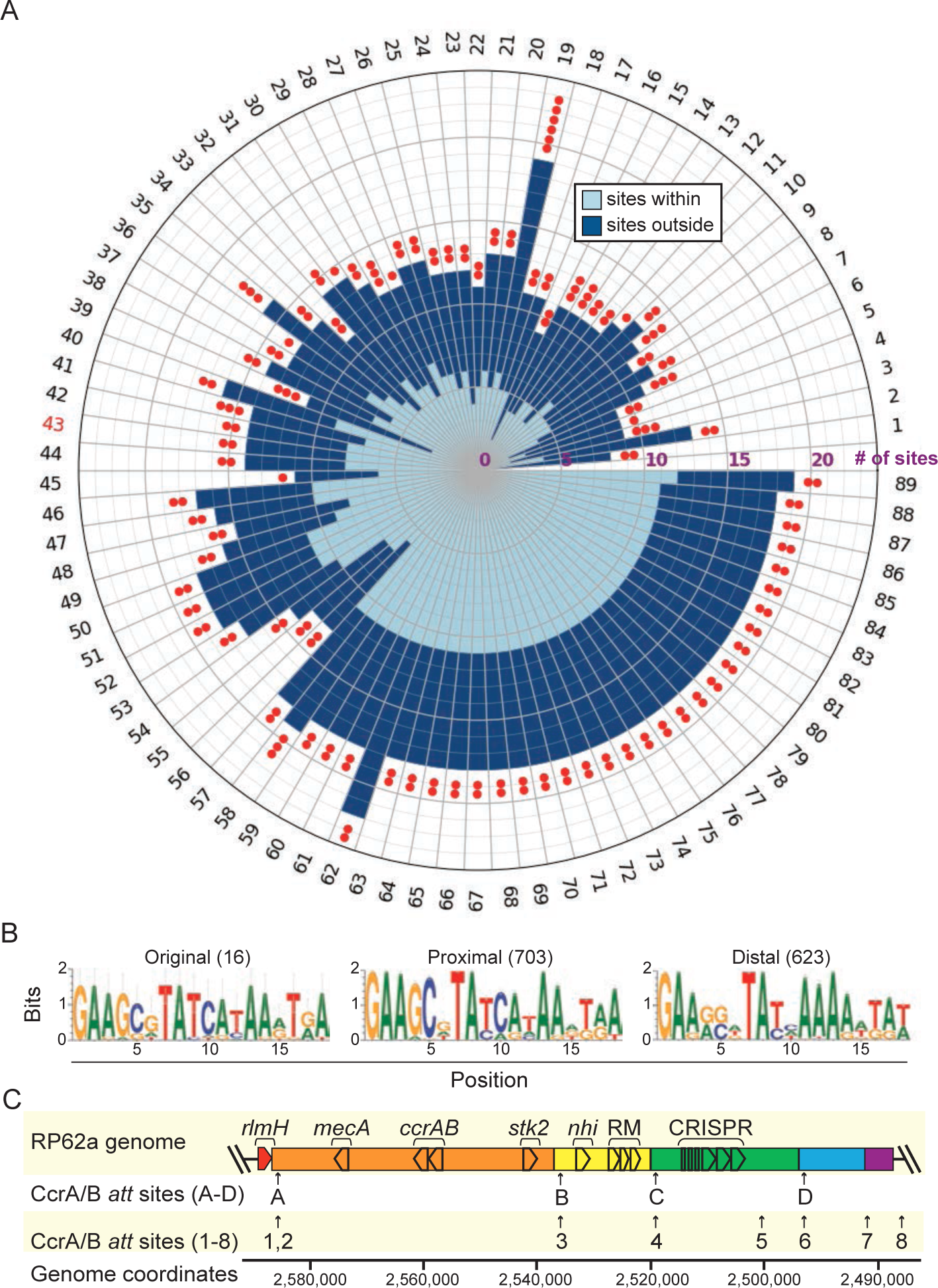
Accompanies Figure 3. Predicted CcrAB *att* sites occur distal to the SCC*mec*/accessory region. (A) Numbers of CcrAB *att* consensus sites within 89 *S. epidermidis* genomes are shown as defined by a motif compiled from 16 experimentally validated sites. Numbers of sites that occur proximal (light blue) and distal (dark blue) to the SCC*mec* accessory region are indicated as a stacked bar graph plotted on a polar axis. Numbers of red dots indicate the number of putative distal cassettes (defined by a segment <150 kb flanked by *att* sites on the same strand). Tip labels correspond to genome number, and the label for *S. epidermidis* RP62a appears in red. (B) Sequence logos built from the 16 validated *att* sites (left) 703 *att* sites proximal (center) and 623 *att* sites distal (right) to SCC*mec*. (C) Illustration of the SCC*mec* accessory region of *S. epidermidis* RP62a showing positions of the eight proximal *att* sites that were detected. Additional putative cassettes are colored in blue and purple.

**Extended data Figure 6.**
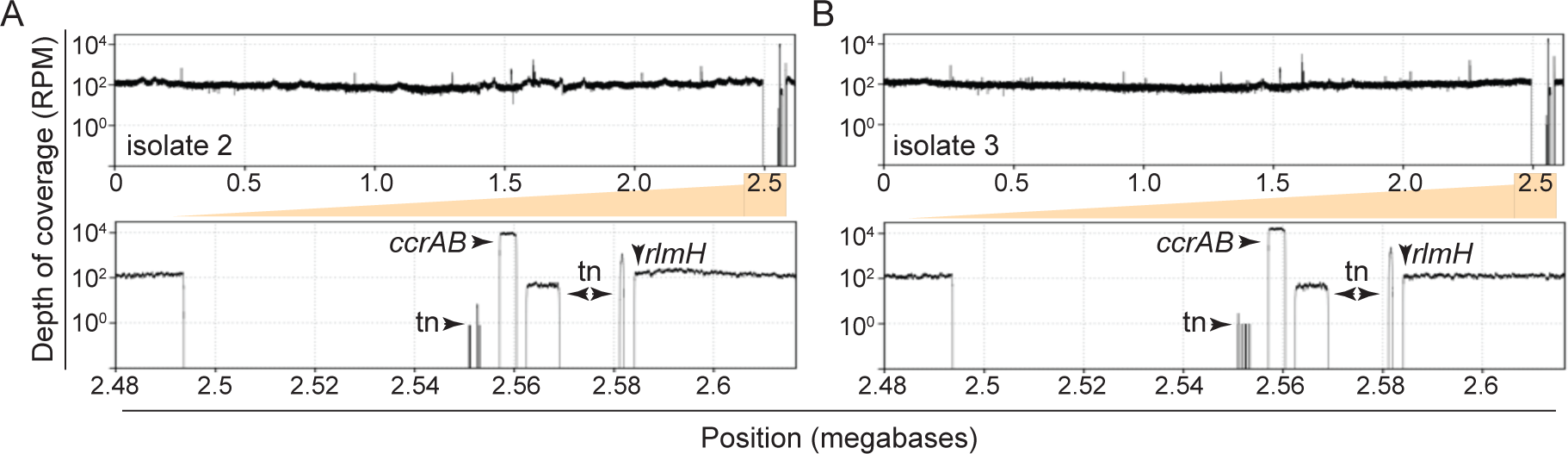
Accompanies Figure 3. Mapped Illumina sequencing reads for two additional *S. epidermidis* RP62a 1′cassette isolates. The reads from representative evolved 1′cassette isolates 2 (A) and 3 (B) were mapped back onto an assembly of the wild-type ancestral genome. The plots show depth of coverage in reads per million (RPM) across the genomes. The bottom plots zoom into the sole genomic segment in the isolates with missing coverage. Arrows indicate read coverage originating from the plasmid-encoded *ccrAB* operon and transposable elements (tn) represented in other regions of the genome.

## References

1. Doron, S. et al. Systematic discovery of antiphage defense systems in the microbial pangenome. Science. 359, eaar4120 (2018).

2. Gao, L. et al. Diverse enzymatic activities mediate antiviral immunity in prokaryotes. Science. 369, 1077–1084 (2020).

3. Vassallo, C. N., Doering, C. R., Littlehale, M. L., Teodoro, G. I. C. & Laub, M. T. A functional selection reveals previously undetected anti-phage defence systems in the E. coli pangenome. Nat. Microbiol. 7, 1568–1579 (2022).

4. Takahashi, T. & Kikuchi, N. Phylogenetic relationships of 38 taxa of the genus Staphylococcus based on 16s rRNA gene sequence analysis. Int. J. Syst. Bacteriol. 49, 725–728 (1999).

5. Becker, K., Heilmann, C. & Peters, G. Coagulase-negative staphylococci. Clin. Microbiol. Rev. 27, 870–926 (2014).

6. Grice, E. A. & Segre, J. A. The skin microbiome. Nat. Rev. Microbiol. 9, 244–253 (2011).

7. Chalmers, S. J. & Wylam, M. E. Methicillin-Resistant Staphylococcus aureus Infection and Treatment Options. in Methicillin-Resistant Staphylococcus Aureus (MRSA) Protocols (ed. Ji, Y.) vol. 2069 (Humana, 2020).

8. Otto, M. Staphylococcus epidermidis—the’accidental’pathogen. Nat. Rev. Microbiol. 7, 555–567 (2009).

9. Otto, M. Coagulase-negative staphylococci as reservoirs of genes facilitating MRSA infection: Staphylococcal commensal species such as Staphylococcus epidermidis are being recognized as important sources of genes promoting MRSA colonization and virulence. BioEssays 35, 4–11 (2013).

10. Ito, T., Katayama, Y. & Hiramatsu, K. Cloning and nucleotide sequence determination of the entire mec DNA of pre-methicillin-resistant Staphylococcus aureus N315. Antimicrob. Agents Chemother. 43, 1449–1458 (1999).

11. Tetsuo Yamaguchi, Ono, D. & Sato, A. Staphylococcal Cassette Chromosome mec (SCCmec) Analysis of MRSA. in Methods in Molecular Biology vol. 2069 59–78 (2020).

12. Centers for Disease Control and Prevention. Antibiotic resistance threats in the United States 2019. http://www.cdc.gov/DrugResistance/Biggest-Threats. (2019) doi:http://www.cdc.gov/DrugResistance/Biggest-Threats.

13. De Oliveira, D. M. P. et al. Antimicrobial Resistance in ESKAPE Pathogens. Clin. Microbiol. Rev. 33, e00181–19 (2020).

14. Centers for Disease Control and Prevention. COVID-19 U.S. Impact on Antimicrobial Resistance. (2022).

15. Lee, J. Y. H. et al. Global spread of three multidrug-resistant lineages of Staphylococcus epidermidis. Nat. Microbiol. 3, 1175–1185 (2018).

16. Hatoum-Aslan, A. The phages of staphylococci: critical catalysts in health and disease. Trends Microbiol. **May** 21, 1–13 (2021).

17. Strathdee, S. A., Hatfull, G. F., Mutalik, V. K. & Schooley, R. T. Phage therapy: From biological mechanisms to future directions. Cell 186, 17–31 (2023).

18. Doub, J. B. et al. Experience Using Adjuvant Bacteriophage Therapy for the Treatment of 10 Recalcitrant Periprosthetic Joint Infections: A Case Series. Clin. Infect. Dis. 76, 1463– 1466 (2023).

19. Aslam, S. et al. Lessons learned from the first 10 consecutive cases of intravenous bacteriophage therapy to treat multidrug-resistant bacterial infections at a single center in the United States. Open Forum Infect. Dis. 7, (2020).

20. Sjöström, J.-E., Löfdahl, S. & Philipson, L. Biological Characteristics of a Type I Restriction-Modification System in Staphylococcus aureus. J. Bacteriol. 133, 1144–1149 (1978).

21. Marraffini, L. A. & Sontheimer, E. J. CRISPR interference limits horizontal gene transfer in staphylococci by targeting DNA. Science. 322, 1843–1845 (2008).

22. Depardieu, F. et al. A Eukaryotic-like Serine / Threonine Kinase Protects Staphylococci against Phages. Cell Host Microbe 20, 471–481 (2016).

23. Bari, S. M. N. et al. A unique mode of nucleic acid immunity performed by a multifunctional bacterial enzyme. Cell Host Microbe 30, 570–582 (2022).

24. Millman, A. et al. An expanded arsenal of immune systems that protect bacteria from phages. Cell Host Microbe 30, 1556–1569 (2022).

25. Rousset, F. et al. Phages and their satellites encode hotspots of antiviral systems. Cell Host Microbe 30, 740–753 (2022).

26. Abby, S. S., Néron, B., Ménager, H., Touchon, M. & Rocha, E. P. C. MacSyFinder: A Program to Mine Genomes for Molecular Systems with an Application to CRISPR-Cas Systems. PLoS One 9, e110726 (2014).

27. Tesson, F. et al. Systematic and quantitative view of the antiviral arsenal of prokaryotes. Nat. Commun. 13, 1–10 (2022).

28. Uehara, Y. Current Status of Staphylococcal Cassette Chromosome mec (SCCmec). Antibiotics 11, 1–12 (2022).

29. Liu, J. et al. Staphylococcal chromosomal cassettes mec (SCC mec): A mobile genetic element in methicillin-resistant Staphylococcus aureus. Microb. Pathog. 101, 56–67 (2016).

30. Wang, L. & Archer, G. L. Roles of CcrA and CcrB in Excision and Integration of Staphylococcal Cassette Chromosome mec, a Staphylococcus aureus Genomic Island. J. Bacteriol. 192, 3204–3212 (2010).

31. Misiura, A. et al. Roles of two large serine recombinases in mobilizing the methicillin-resistance cassette SCCmec. Mol. Microbiol. 88, 1218–1229 (2013).

32. O’Connor, A. M. et al. Significant Enrichment and Diversity of the Staphylococcal Arginine Catabolic Mobile Element ACME in Staphylococcus epidermidis Isolates From Subgingival Peri-implantitis Sites and Periodontal Pockets. Front. Microbiol. 9, 1–15 (2018).

33. Holt, D. C. et al. A Very Early-Branching Staphylococcus aureus Lineage Lacking the Carotenoid Pigment Staphyloxanthin. Genome Biol. Evol. 3, 881–895 (2011).

34. Tong, S. Y. C. et al. Novel staphylococcal species that form part of a Staphylococcus aureus-related complex: the non-pigmented Staphylococcus argenteus sp. nov. and the non-human primate-associated Staphylococcus schweitzeri sp. nov. Int. J. Syst. Evol. Microbiol. 65, 15–22 (2015).

35. Golding, G. R. et al. Whole-Genome Sequence of Livestock-Associated ST398 Methicillin-Resistant Staphylococcus aureus Isolated from Humans in Canada. J. Bacter 194, 6627–6628 (2012).

36. Maniv, I., Jiang, W., Bikard, D. & Marraffini, L. A. Impact of different target sequences on type III CRISPR-Cas immunity. J. Bacteriol. 198, 941–950 (2016).

37. Diep, B. A. et al. The Arginine Catabolic Mobile Element and Staphylococcal Chromosomal Cassette mec Linkage: Convergence of Virulence and Resistance in the USA300 Clone of Methicillin-Resistant Staphylococcus aureus. J. Infect. Dis. 197, 1523– 30 (2008).

38. Connor, A. M. O., Mcmanus, B. A. & Coleman, D. C. First description of novel arginine catabolic mobile elements (ACMEs) types IV and V harboring a kdp operon in Staphylococcus epidermidis characterized by whole genome sequencing. Infect. Genet. Evol. 61, 60–66 (2018).

39. Almebairik, N. et al. Genomic Stability of Composite SCC mec ACME and COMER-Like Genetic Elements in Staphylococcus epidermidis Correlates With Rate of Excision. Front. Microbiol. 11, 1–12 (2020).

40. Scharn, C. R., Tenover, F. C. & Goering, R. V. Transduction of staphylococcal cassette chromosome mec elements between strains of Staphylococcus aureus. Antimicrob. Agents Chemother. 57, 5233–5238 (2013).

41. Mašlaňová, I. et al. Bacteriophages of Staphylococcus aureus efficiently package various bacterial genes and mobile genetic elements including SCCmec with different frequencies. Environ. Microbiol. Rep. 5, 66–73 (2013).

42. Chlebowicz, M. A. et al. The Staphylococcal Cassette Chromosome mec type V from Staphylococcus aureus ST398 is packaged into bacteriophage capsids. Int. J. Med. Microbiol. 304, 764–774 (2014).

43. Ray, M. D., Boundy, S. & Archer, G. L. Transfer of the methicillin resistance genomic island among staphylococci by conjugation. Mol. Microbiol. 100, 675–685 (2016).

44. Maree, M. et al. Natural transformation allows transfer of SCCmec-mediated methicillin resistance in Staphylococcus aureus biofilms. Nat. Commun. 13, 1–14 (2022).

45. Fillol-Salom, A. et al. Bacteriophages benefit from mobilizing pathogenicity islands encoding immune systems against competitors. Cell 185, 3248–3262 (2022).

46. LeGault, K. N., Barth, Z. K., DePaola, P. & Seed, K. D. A phage parasite deploys a nicking nuclease effector to inhibit viral host replication. Nucleic Acids Res. 50, 8401– 8417 (2022).

47. Pinilla-Redondo, R. et al. CRISPR-Cas systems are widespread accessory elements across bacterial and archaeal plasmids. Nucleic Acids Res. 50, 4315–4328 (2022).

48. Benler, S. et al. Cargo Genes of Tn 7-Like Transposons Comprise an Enormous Diversity of Defense Systems, Mobile Genetic Elements, and Antibiotic Resistance Genes. MBio 12, e02938–21 (2021).

49. LeGault, K. N. et al. Temporal Shifts in Antibiotic Resistance Elements Govern Phage-Pathogen Conflicts. Science. 373, 1–29 (2021).

50. Dedrick, R. M. et al. Prophage-mediated defence against viral attack and viral counter-defence. Nat. Microbiol. 2, 1–13 (2017).

51. Owen, S. V et al. Prophages encode phage-defense systems with cognate self-immunity. Cell Host Microbe 29, 1–14 (2021).

52. Berger, B., Waterman, M. S. & Yu, Y. W. Levenshtein Distance, Sequence Comparison and Biological Database Search. IEEE Trans Inf Theory 67, 3287–3294 (2021).

53. Potter, S. C., et al. HMMER web server: 2018 update. Nucleic Acids Res. 46, W200– W204 (2018).

54. Christensen, G. D., Baddour, L. M. & Simpson, W. A. Phenotypic variation of Staphylococcus epidermidis slime production in vitro and in vivo. Infect. Immun. 55, 2870–2877 (1987).

55. Nair, D. et al. Whole-Genome Sequencing of Staphylococcus aureus Strain RN4220, a Key Laboratory Strain Used in Virulence Research, Identifies Mutations That Affect Not Only Virulence Factors but Also the Fitness of the Strain. J. Bacteriol. 193, 2332–2335 (2011).

56. Kwong, S. M., Ramsay, J. P., Jensen, S. O. & Firth, N. Replication of Staphylococcal Resistance Plasmids. Front. Microbiol. 8, 1–16 (2017).

57. Gibson, D. G. et al. Enzymatic assembly of DNA molecules up to several hundred kilobases. Nat Meth 6, 343–345 (2009).

58. Monk, I. R., Shah, I. M., Xu, M., Tan, M. & Foster, T. J. Transforming the Untransformable: Application of Direct Transformation To Manipulate Genetically Staphylococcus aureus and Staphylococcus epidermidis. MBio 3, 1–11 (2012).

59. Cater, K. et al. A Novel Staphylococcus Podophage Encodes a Unique Lysin with Unusual Modular Design. 2, e00040–17 (2017).

60. Vandersteegen, K. et al. Microbiological and Molecular Assessment of Bacteriophage ISP for the Control of Staphylococcus aureus. PLoS One 6, e24418 (2011).

61. Freeman, M. E. et al. Complete Genome Sequences of Staphylococcus epidermidis Myophages Quidividi, Terranova, and Twillingate. Microbiol. Resour. Announc. 8, e00598–19 (2019).

62. Jiang, W. et al. Dealing with the Evolutionary Downside of CRISPR Immunity: Bacteria and Beneficial Plasmids. PLoS Genet. 9, e1003844 (2013).

